# Haplotype specific analyses in the phased genomes era: the case of apple cv ‘Golden Delicious’

**DOI:** 10.64898/2026.01.28.702214

**Authors:** Luca Bianco, Nicola Busatto, Mirko Moser, Diego Micheletti, Lorenzo Spina, Michela Troggio, Stefano Piazza, Fabrizio Costa, Paolo Fontana

## Abstract

Apple is one of the major cultivated fruit crops in temperate regions. To better support breeding programs and facilitate the development of improved cultivars, we generated a new haplotype-resolved version of the Golden Delicious genome, one of the major founders of many modern apple lineages. The assembly features the separation of the two haplotypes, with a total size of 647.3 Mb and 649.2 Mb, respectively. The phasing was accurately validated with 10,321 curated SNPs. Telomere-to-telomere continuity was verified by the analysis of telomeric sequence composition at the end of each chromosome. Gene prediction identified a total of 45,116 genes in haplotype 1 and 45,063 genes in haplotype 2. A pangenome analysis employing 6 haplotype resolved genomes identified both common and unique gene families. The availability of a phased genome enabled the assessment of genome-wide allelic specific expression. Our case study, focusing on *Md-PG1* (a key regulator of fruit softening), revealed that the allelic form on haplotype 2 (GDH2-10g24673) was the dominant contributor to total gene expression. In addition, the phased genome also showed specific miRNA chromosomal distribution patterns, as well as a distinct methylation profile. Altogether, these genomic resources provide new insights into the allelic regulation of key agronomic traits and represent a valuable tool to accelerate apple breeding.

## Introduction

Life science disciplines have greatly benefited from the application of sequencing technologies, which have enabled the deciphering of the DNA-nucleotide alphabet in many living organisms. From the first pioneering Sanger method, a series of novel and improved technologies has been developed, contributing to a substantial advancement in unraveling the genome structure of sequenced organisms. The advent of long read sequencing, and its improvement in terms of read length and base-level accuracy, has opened new frontiers allowing the assembly of chromosome-scale haplotype-resolved genomes. Traditionally, genome assemblies represented a consensus between the two (in case of diploid genomes) or more haplotypes, collapsing slightly different sequences, losing, therefore, haplotype-specific variations. The latest sequencing technologies now enable the resolution of haplotype-specific alleles, structural variants, and SNPs that are typically merged in consensus genome assemblies. The information encoded in haplotypes is fundamental to untangle the genetic structure and diversity of a population, essential prerequisites for haplotype-based breeding (HBB), a promising approach for improving crop varieties through the identification of superior haplotypes to integrate into breeding programs (Sivabharathi *et al*., 2024; Peleman *et al*., 2003).

HBB is a promising but still emerging technique in crop breeding: its foundation involves creating haplotype maps for a crop population. This is achieved by sequencing and assembling the phased genomes of several individuals. This initial map allows breeders to identify common and rare haplotypes, which can then be used alongside genotyping data from a larger population to uncover the genetic diversity (Bhat *et al*., 2021).

Complementary pangenome analyses further enable the detection of structural variants, introgressed haplotypes, and accession-specific genes, including those associated with pest and pathogen resistance (Zhao *et al*., 2018; Garg *et al*., 2022). When combined with genome-wide association studies (GWAS), haplotype-level analyses provide increased power to detect loci underlying complex traits, as they capture the combined effect of linked variants (Qian *et al*., 2017).

Apple (*Malus x domestica*), one of the most economically important cultivated fruit crops, has recently benefited from haplotype-resolved genomic approaches (Sun *et al*., 2020; Khan *et al*., 2022; Cai *et al*., 2024; Švara *et al*., 2024; *Mansfeld et al*., 2023). Pangenome analysis of phased genomes in the cultivar ‘Gala’ together with two related accessions (*M. sieversii* and *M. sylvestris*), revealed that its genome has a 23% hybrid origin. The inclusion of an additional 91 accessions provided a better understanding of the genetic basis of domestication and the improvement of important fruit quality traits (Sun *et al*., 2020). In the same species, haplotype-resolved genome assembly coupled with pangenome analysis revealed the proportion of genes gained and lost during domestication, especially in resistance genes analogs (RGAs) (Su *et al*., 2024). Similar approaches also allowed the identification of 376 RGAs possibly originated by duplication events in avocado (Yang *et al*., 2024). In kiwifruit (*Actinidia chinesis*), haplotype-phased genome sequencing revealed the level of heterozygosity between the two homologous chromosomes of the two haplotypes, offering valuable perspectives for future breeding programs (Yue *et al*., 2024).

Furthermore, RNA-Seq experiments exploiting the haplotype resolved genomes can identify allele-specific expression (ASE) patterns and dominant or recessive alleles that differently contribute to the phenotype (Hasing *et al*., 2020; Seo *et al*., 2016; Zhou *et al*., 2020). Haplotype-resolved genome assembling also opened the possibility to capture and characterize allelic differences across germplasm collections or wild species, which resulted in highly efficient breeding processes (Bhat *et al*., 2021). ASE refers to the preferential expression of one allele in a heterozygous individual (Gaur *et al*., 2013), and can be influenced by genetic differences, epigenetic modifications, and environmental factors (St Pierre *et al*., 2022). Analysis of allelic expression imbalance can therefore provide insights into interactions between alleles of subgenomes, considered one of the mechanisms underlying heterosis (Goff and Zhang, 2013; Shao *et al*., 2019). In pear, the availability of a haplotype resolved genome enabled the characterization of ASE bias in the parental haplotype of a specific hybrid, and allelic specific expression patterns were found in genes involved in important quality traits, such as sugar accumulation (Li *et al*., 2024). In poplar, a haplotype-resolved genome coupled with ASE analysis revealed an unbalanced expression in 37% of the alleles (Shi *et al*., 2024).

Here, we present a haplotype-resolved genome assembly of the cultivar ‘Golden Delicious’ (*Malus x domestica*), the reference cultivar for apple research, and a recurrent parental line in many modern breeding initiatives. We analyze the haplotypes of this cultivar to provide information on the genome structure also in comparison with other five haplotype resolved apple genomes, demonstrating the applicability of these resources through two advanced analyses: genome-wide allele specific expression and haplotype-resolved DNA methylation.

## Materials and methods

### Whole genome sequencing, assembling and sequence annotation

For DNA extraction, young leaves were collected in spring from a ‘Golden Delicious’ clone B tree grown at the experimental orchard of the Fondazione Edmund Mach, located in the Northern part of Italy. High-molecular-weight genomic DNA, intended for both PacBio HiFi and Omni-C sequencing strategies, was extracted using the NucleoBond HMW DNA Kit (Macherey-Nagel, Germany) according to the manufacturer’s protocol. Library preparation and sequencing for the PacBio HiFi strategy were performed at Novogene (Cambridge, UK) using the PacBio Sequel II platform based on single-molecule real-time (SMRT) sequencing technology. Library preparation and sequencing for the Omni-C strategy were carried out at SciLifeLab (Stockholm, SE).

PacBio HiFi reads together with Omni-C reads were assembled using the hifiasm assembler v.0.19.0 (Cheng *et al*., 2021). The resulting contigs were scaffolded into chromosomes using the software ALLMAPS v.1.3.3 integrating information retrieved from the dense integrated genetic map (iGLMap) available for apple (Di Pierro E, *et al*., 2016). Phasing accuracy was verified by aligning, for both assembled haplotypes, the probes (i.e. flanking regions) of a core set of 10,321 Golden Delicious phased single nucleotide polymorphism (SNP) markers included in the Infinium Apple 20K array (Bianco *et al*., 2016, Howard *et al*., 2021). Only single mapping probes, with 98% or higher identity and 98% or higher coverage of the full sequence, were retained for downstream analysis. The comparison of the phase of both sequences and mapped SNPs was carried out with a custom python script (available upon request).

The terminal regions of each chromosome (1kbps) were analyzed with a custom-made python script to identify the usual plant telomeric repeat 3’-TTTAGGG-5’.

BUSCO v.5 software (Manni et al., 2021) was used to assess the gene-space completeness of the two haplotypes and the LTR Assembly Index (LAI) was computed using the software LTR_retriever v.3.0.1 (Ou et al., 2018).

Sequence collinearity between the ‘Golden Delicious’ phased genome assembled in this paper and the ‘Golden Delicious’ doubled-haploid GDDH13 genome (Daccord et al., 2017) was assessed with dgenies v. 1.5.0 (Cabanettes and Klopp, 2018), while possible structural variants across the two haplotypes and the GDDH13 genome were assessed with the software Syri v.1.7.0 (Goel et al., 2019).

### Gene prediction and functional annotation

Repeat masking was performed using the *Rosaceae* library downloaded from RpetDB v2. Coding gene prediction was performed separately for each haplotype using Augustus v3.4 (Stanke *et al*., 2006) and the MAKER v.3.01 pipeline (Canterel *et al*., 2008), comprising GeneMark-ES, Augustus and EVidenceModeler, trained with RNASeq downloaded from SRA (SRX10603975, SRX10603976, SRX10603977, SRX10603978, SRX10808467, SRX10808468, SRX10808469, SRX10808470, SRX2697721, SRX2697722, SRX2697723, SRX2697724, SRX3408565, SRX3408566, SRX3408567, SRX3408568, SRX3408569, SRX3408570, SRX3408571, SRX3408576, SRX3408577, SRX3408578, SRX3420319, SRX3420330, SRX3420397, SRX4216806, SRX4216807) and proteins downloaded from NCBI RefSeq belonging to *Malus x domestica*. The two predictions were merged according to the results of the GeneValidator v.2.1.12 tool (Drăgan *et al*., 2016), which retained only the best predictions for every gene site. Gene prediction data quality was evaluated with BUSCO tool v 5.5.0 (Manni *et al*., 2021). tRNAs were predicted with tRNAscan-SE v2.0.6 (Chan *et al*., 2019). Predicted genes were functionally annotated with Gene Ontology (GO) terms using Argot2 (Falda *et al*., 2012), and results were filtered using FunTaxIS-lite (Bianca *et al*., 2023).

### Comparative genomics

Five haplotype-resolved genomes were collected for comparison with the newly assembled ‘Golden Delicious’ genome: *Malus x domestica* cv. ‘Gala’ (*Sun et al*., 2020), *Malus x domestica* cv. ‘Honeycrisp’ (*Khan et al*., 2022), *Malus x domestica* cv. ‘Fuji’ (*Cai et al*., 2024), *Malus x domestica* cv. ‘Antonovka’ (*Švara et al*., 2024), and *Malus fusca* (*Mansfeld et al*., 2023). Orthofinder with default parameters was used to identify homology relationships among the considered accessions. The pangenome graph was built using the Cactus pipeline with --clip 0 --filter 0 parameters, setting haplotype 1 of ‘Golden Delicious’ as reference. Collinearity analysis was performed with the MCscan (Python version) (*Tang et al*., 2024) pipeline using default parameters.

### Haplotype resolved allelic specific expression (ASE)

Allelic specific expression was analysed through a re-analysis of the whole-genome expression performed on apple samples collected during the ripening progression (Busatto *et al*., 2021).

Libraries were clustered on a cBot Cluster using the TruSeq SR Cluster Kit v3 and sequenced on the Illumina HiSeq2000 platform with the TruSeq SBS Kit v3. Sequence data are available at the GEO database (Accession N.:GSE182822; Busatto *et al*., 2021) and SRA (PRJNA1399370).

Sequencing reads were aligned to both haplotypes of the newly assembled ‘Golden Delicious’ genome using HISAT2 to identify reads aligning on more than one locus. The MCscan results were further employed to identify specific gene alleles on each haplotype, and a custom Python script was developed to count reads that uniquely aligned to a single allele. This process ensured that each read was unambiguously assigned to one of the two alleles, creating a reliable count table. The resulting read count table was used as input for DESeq2 (*Love et al*., *2014*) to identify differentially expressed alleles. An allele was considered as differentially expressed when its p-value was less than or equal to 0.05. This statistical analysis allowed to highlight significant differences in expression levels between the two haplotypes.

In addition, a traditional approach was used for counting reads per gene ID on each haplotype using the tool featureCounts (*Liao et al*., *2014*), providing the haplotype-specific General Feature Format (gff) file with annotations. Differential expression analysis was carried out within each haplotype between the two considered timepoints of the maturation-to-ripening experiment. The two independent sets of differentially expressed genes were then compared both with the set of transcripts showing haplotype specific expression highlighting DEGs whose expression is predominantly derived from one or the other haplotype.

### Allele-specific microRNA analysis

Sequences of the *Malus x domestica* mature miRNAs and pre-miRNA (hairpin) were retrieved from miRBase (version R22.1, https://mirbase.org/, Griffiths-Jones et al., 2008). Mature sequences were aligned against each haplotype of the newly assembled ‘Golden Delicious’ genome separately using bowtie (version 1.3.1; Langmead et al., 2009) with default parameters except for parameter v=0. Hairpin sequences were aligned using blastn (Altschul et al., 1990) with –evalue=1E-7 –num_alignments 5. Regions where mature and hairpin sequences aligned with overlap were considered as complete loci, whereas regions with either only mature or hairpin alignments were indicated as possible variants. Comparative analysis of the miRNA loci was then performed between the two haplotypes, classifying loci as common, when present in both haplotypes; variable, when belonging to a miRNA family with different number of loci between the two haplotypes and unique, when present only in one or the other haplotype.

### Allele-specific methylation analysis

Haplotype specific methylation was analysed by sequencing the DNA extracted from young leaves of ‘Golden Delicious’ using Oxford Nanopore Technology (ONT). DNA was extracted with the NucleoSpin Plant II Midi (Macherey-Nagel, Germany) following the manufacturer’s protocol. Libraries were prepared using the Ligation Sequencing Kit V14 (SQK-LSK114) following the manufacturer’s protocol, and loaded onto a R10.4 flow cell (FLO-PRO114M). Base calling was performed using the super-accurate model in MinKNow v. 24.06.10. Cytosine modifications (5mC and 5hmC) were identified with the software Dorado v.7.4.12, using the model dna_r10.4.1_e8.2_400bps_sup@v4.3.0_5mC_5hmC@v1.

ONT reads were aligned to the final version of the haplotype resolved genome of ‘Golden Delicious’ using MINIMAP2 (Li, 2018) with the “map-ont” preset. Methylation pattern analysis was performed after removing the reads with mapping quality <10 and keeping only primary alignments, using the pileup and summary functions of modkit (https://github.com/nanoporetech/modkit). Differentially methylated genomic regions between haplotypes were identified using Methylartist (Cheetham et al., 2022). The differential methylation value in collinear genes was measured using the – Log2FoldChange calculated as the ratio between the methylation fraction in Haplotype 1 and Haplotype 2.

## Results and Discussion

### Golden Delicious genome assembly

PacBio sequencing of the selected ‘Golden Delicious’ sample produced 2.15M HiFi reads for a total of 41,81Gb of data for an estimated ∼56x coverage (based on the haploid estimated genome size). The N50 of the HiFi reads was 19,557 bps. Regarding Omni-C, a total of 471M unique 151bp paired-end reads were produced, resulting in ∼90x coverage. The ‘Golden Delicious’ genome sequence was assembled into two sets of 17 chromosomes (haplomes) using hifiasm and ALLMAPS resulting in a total genome size of Haplotype 1 of 647.3 Mb with a N50 of 36.8 Mb, while Haplotype 2 was slightly bigger with 649.2 Mb and N50 of 35.9 Mb. Marey plots (Supplementary Figure S1) generated by ALLMAPS were used to assess the correlation between the iGLMap genetic map (Di Pierro E, et al., 2016) and the assembled sequences, highlighting a strong correlation between the physical and genetic coordinates (correlation > 0.99 for all chromosomes; see Supplementary Figures 1-17 for Marey plots).

Long terminal repeat (LTR) assembly analysis resulted in a LAI index of 18.93 for Haplotype 1 and 18.65 for Haplotype 2, empirically categorizing both haplotypes as ‘reference-quality’ (i.e. LAI between 10 and 20) assemblies of the repetitive and intergenic space (Ou *et al*., 2018).

Analysis of the terminal regions of each chromosome identified 30 of 34 telomeric repeats in Haplotype 1 and 28 of 34 repeats in Haplotype 2, showing that most of the chromosomes were to near telomere-to-telomere resolution (T2T).

BUSCO analysis on the eudicots dataset (n=2,326) of both ‘Golden Delicious’ haplotypes together evidenced a high gene space completeness, with 99.0% of complete genes recovered and only 0.6% of fragmented and 0.4% of missing.

To further validate the accuracy of the haplotype resolution of the newly assembled Golden Delicious genome, probes of 10,321 phased SNPs were aligned on the genome. Out of these, 9,735 SNPs passed the filters on Haplotype 1 and 9,774 SNPs on Haplotype 2. Among these, 9,474 SNPs were present in both haplotypes, of which 4,298 were informative (heterozygous AB), while the remaining 5,176 SNPs were present in both haplotypes in homozygous state (AA/BB) and therefore not informative for phasing. Haplotype assessment confirmed that the phase of the set of polymorphic SNPs corresponded to the phase of the assembled sequences, with the exception of a single SNP (SNP_FB_1103570), located in chr8 (at position 27,199,459 in Haplotype1, 27,728,092 in Haplotype2, see supplementary material). Based on phasing and pedigree information, Haplotype 1 was assigned as the chromosome set inherited from Grimes Golden, which is the only known and verified parent of Golden Delicious. Structural variant analysis using the software Syri confirmed previously reported rearrangements (e.g. the chromosome 1 inversion in GDDH13) and additionally identified two consecutive inversions in the central region of Ch6 of Haplotype 1, located at 10,9 - 13,2 Mb and 13,2 - 16,4 Mb, respectively (Supplementary Figure S2), affecting 34 genes and 55 genes respectively (Supplementary Table S1 and S2).

Overall, these analyses indicate that the newly assembled ‘Golden Delicious’ haplotypes are highly contiguous, complete and accurately phased.

### Genome annotation

We used a combination of de-novo, RNA-Seq and homology-based methods to target gene structures in the two Golden Delicious haplotypes. Gene prediction found a total of 45,116 CDS with a mean length of 1,236 bp and mean introns per transcripts of 3.9 in the Haplotype 1. In the Haplotype 2, the number of CDS was 45,063 with a mean length of 1,238 bp and mean introns per transcript of 3.9. BUSCO score of the predicted CDS of Haplotype 1 was 99% (completed genes 98.7% and fragmented genes 0.3%), while for Haplotype 2 the score was 98.9% (completed genes 98.6% and fragmented genes 0.3%).

### Pangenome analysis

To investigate genetic diversity in apple, a total of six accessions with available haplotype-resolved genomes were considered for the construction of the pangenome, of which four were *M. domestica* cultivars and two were wild species accessions. The analysis carried out with Orthofinder allowed the investigation of the gene content and functional differences present in apples. According to this analysis, 52,662 predicted gene families were found. Dissecting some of these gene families was observed that the apple pan-genome was represented by 19,532 common gene families with at least one gene present in a single haplotype, and 14,082 common families with genes present on both haplotypes, while the unique gene families (genes present only in one species) were 12,419 (see Fig. 1B). Analysis of the hemizygous and unique gene families revealed that ‘Gala’ exhibited the highest number of hemizygous genes (10,881 across both haplotypes) and the lowest number of unique genes (2,603). In contrast, *M. fusca* showed the highest number of unique genes (13,568) and one of the lowest numbers of hemizygous genes (3,748) (see Table 1).

**Figure 1.**
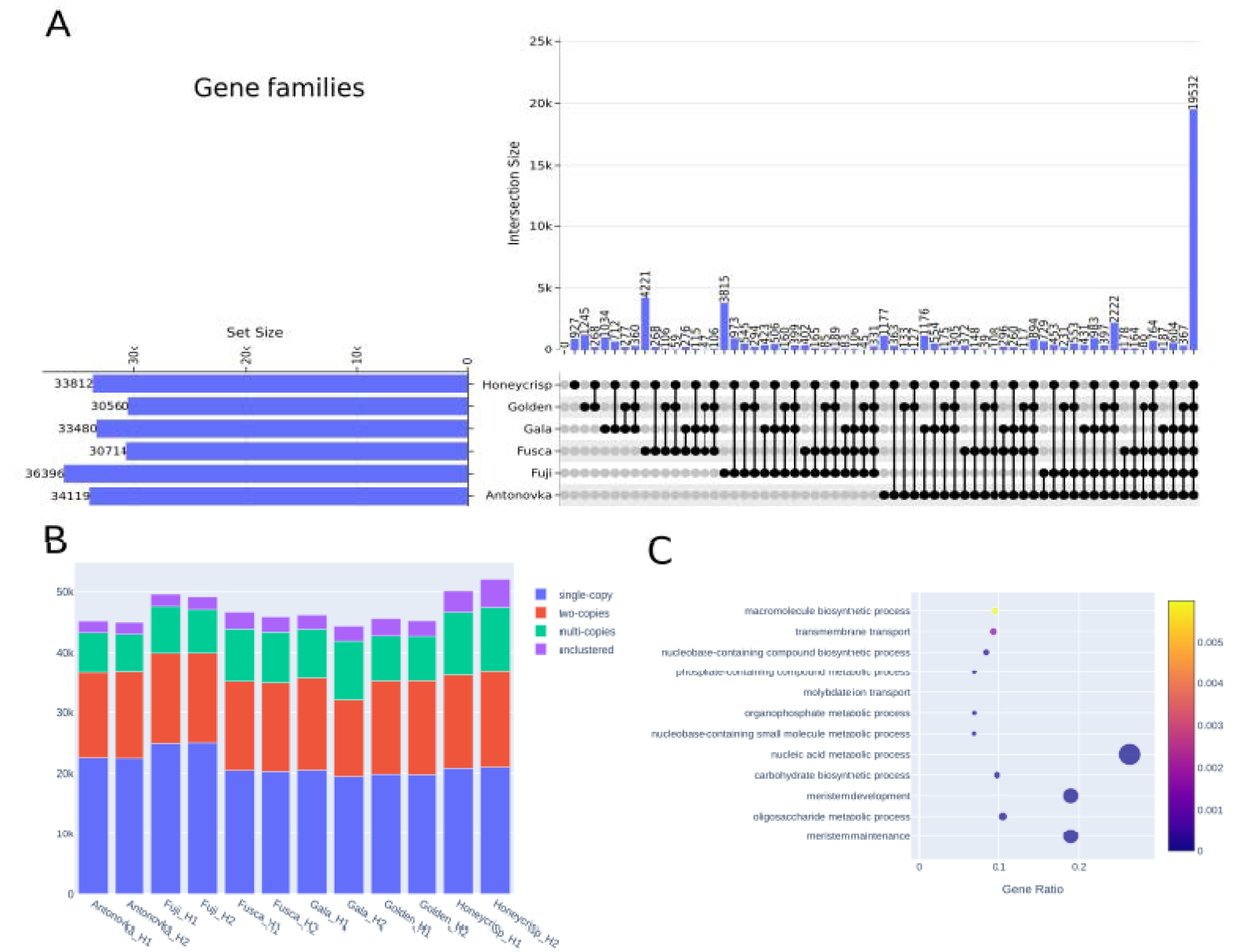
A) Upset plot showing of the shared and unique gene families, as generated with Orthofinder. B) Bar plot summarizes the number of genes per family that are single-copy, two-copies or multi-copy, the unclustered category represents genes that Orthofinder does not assign to any family. C) Scatter plot showing the Gene Ontology functional enrichment of the genes lacking homologues in other apple cultivar or species.

**Table 1.** Hemizygous genes: Genes present on only one haplotype (H1 or H2) within a specific species or cultivar. Unique genes: Genes found exclusively in one of the considered species or cultivar genomes (not present in any of the others). Unique genes: Genes found exclusively in one of the considered species or cultivar genomes (not present in any of the others).

|  | Antonovka |  | Fuji |  | Fusca |  | Gala |  | Golden |  | Honeycrisp |  |
| --- | --- | --- | --- | --- | --- | --- | --- | --- | --- | --- | --- | --- |
|  | H1 | H2 | H1 | H2 | H1 | H2 | H1 | H2 | H1 | H2 | H1 | H2 |
| **Hemizygous genes** | 3422 | 3429 | 2551 | 2482 | 2127 | 1621 | 6408 | 4473 | 1635 | 1652 | 4088 | 5274 |
| **Unique genes** | 1392 | 1304 | 4510 | 4465 | 6857 | 6711 | 1219 | 1384 | 1565 | 1586 | 1560 | 2168 |
| **Unique lost events** | 331 |  | 894 |  | 2222 |  | 764 |  | 604 |  | 367 |  |

The UpSet plot of Fig. 1A reports all the unique and shared ortholog families of the analyzed accessions, while Fig. 1B provides an overview of gene family copy numbers broken down by cultivar and species

Furthermore, GO enrichment analysis of ‘Golden Delicious’ unique genes was performed to investigate its functional peculiarities with respect to the other considered accessions. The resulting GO terms were grouped according to their connection in the GO graph and only the most specific ones were reported. The results clearly showed that the analyzed genes were enriched in macromolecule biosynthetic process, transmembrane transport, nucleobase-containing compound biosynthetic process, phosphate-containing compound metabolic process, molybdate ion transport, organophosphate metabolic process, nucleobase-containing small molecule metabolic process, nucleic acid metabolic process, carbohydrate biosynthetic process, meristem development, and oligosaccharide metabolic process, meristem maintenance (see Fig. 1C).

Pangenome partitioning further highlighted the relative uniqueness of each accession among all the considered genomes for common (Fig. 2A) and unique segments (Figure 2B).

**Figure 2.**
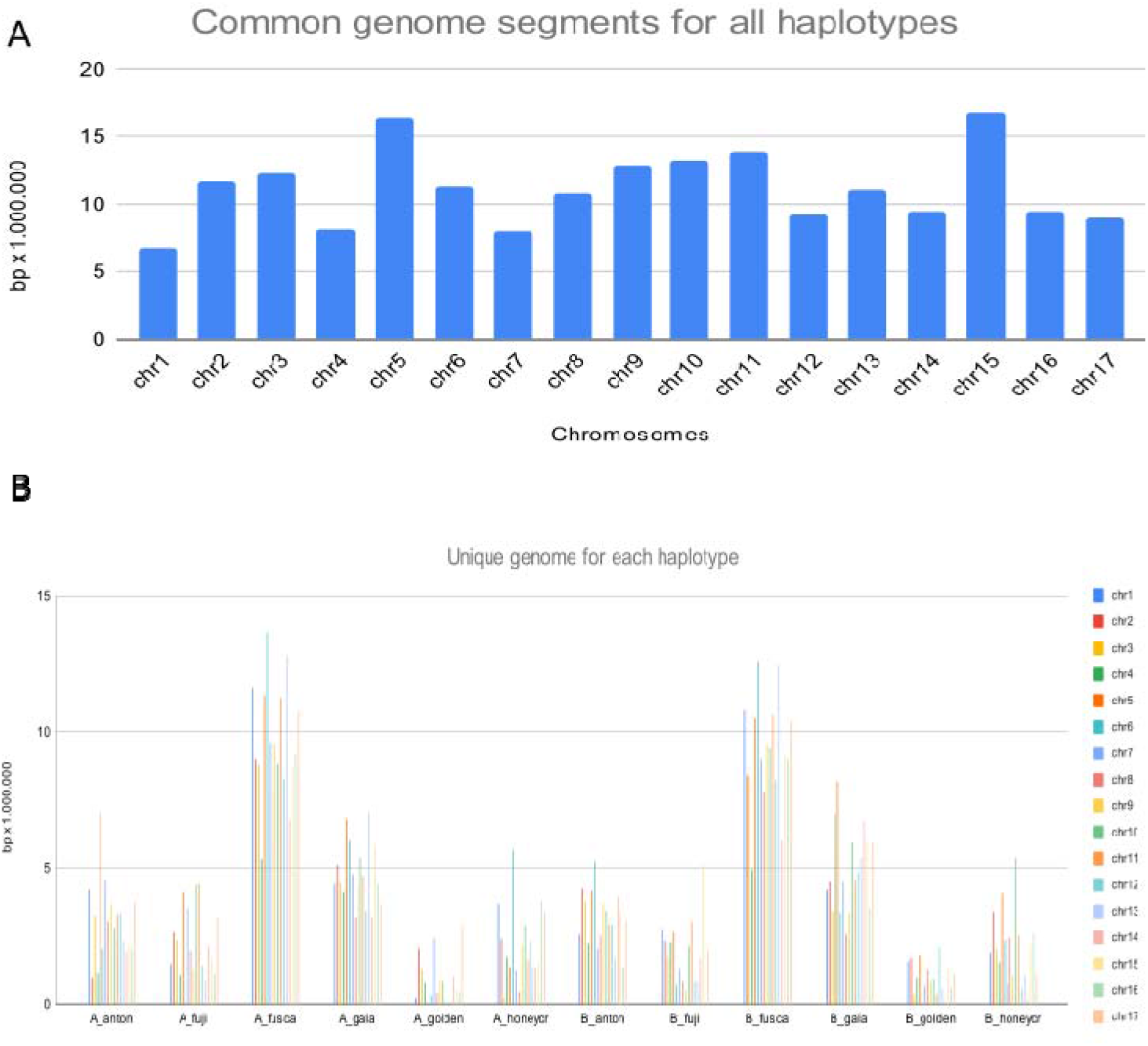
A) Common Genome Segments : This bar plot displays the total length (in base pairs x 1,000,000) of core genome segments—regions shared and in common among all analyzed haplotypes—for each chromosome, as determined by the pangenome analysis. B) Unique Genome Segments : Conversely, this bar plot illustrates the length (in base pairs x1,000,000) of unique genome segments (the accessory/dispensable genome components) contributed by each individual haplotype across the analyzed chromosomes.

As expected, ‘Golden Delicious’ displayed less than 3% of unique genome content per haplotype, the lowest among accessions considered, consistent with its extensive use as a parental genotype in breeding programs. On the other hand, *M. fusca*, a wild species, retained large unique genomic regions: 25.59% for Haplotype 1 and 24.89% for Haplotype 2. Surprisingly, ‘Gala’, a progeny of ‘Golden Delicious’ x ‘Kidd’s Orange Red’, contained a relatively high proportion of unique sequences (12.28% for Haplotype 1 and 14.54% for Haplotype 2), with ∼ 7 Mb region on chromosome 13 (Haplotype 1) not shared with any other cultivar (see Fig. 2B).

### Haplotype-resolved transcriptome analysis reveals H2-specific expression of the polygalacturonase gene MdPG1-3 associated with fruit softening in ‘Golden Delicious’ apple

Traditional consensus genomes merge maternal and paternal haplotypes into a single sequence, thereby losing haplotype-specific expression patterns that are important in functional genomics analyses. With haplotype-resolved genomes, RNA-seq data can be aligned to maternal and paternal genomes separately, allowing detection of allele specific expression (ASE), shedding light on the genetic regulation of both haplotypes. In this regard, this approach represents a significant advancement in the detection of allele-specific expression, as it provides a more straightforward and direct strategy, rather than relying on comparisons between ASE in an F□hybrid and allelic expression in parental lines, an approach that is often not feasible in tree species (Albert et al., 2018). Conversely, in recent studies the association between transcripts and single haplotypes in a T2T phased apple genome has only been briefly addressed, primarily for identifying splicing variants and achieving a more accurate gene annotation (Su *et al*., 2024) or as output of innovative machine-learning modeling (Shi *et al*., 2024). In this context, to highlight the efficacy of our approach, the reads obtained from Busatto *et al*. (2021) were employed, thereby dissecting allele-specific transcriptional regulation, affecting specific aspects of apple ripening, a classic physiological process that can now be interpreted with unprecedented precision through ASE. The Illumina reads capturing transcriptional variation at the climacteric stage of ‘Golden Delicious’ ripening (Busatto et al., 2021) were aligned to the new phased version of the ‘Golden Delicious’ genome, selecting reads that were uniquely mapped to a single haplotype with 100% sequence identity. Among all the pairs of collinear genes showing differential expression identified through this analysis (Supplementary Figure S3), a *polygalacturonase* (*PG*) gene was detected, a key cell wall–modifying enzyme and the primary determinant of the softening pattern in apple (Costa *et al*., 2010). This class of enzymes is responsible for breaking down galacturonic bonds, thereby contributing to the disassembly of the plant cell wall by involving the depolymerization of covalently bound pectin, converting it into a water-soluble form. The main substrate for PGs in the cell wall is homogalacturonan, a type of pectin abundant in the middle lamella. PG activity is closely associated with the later stages of fruit ripening (Alexander and Grierson, 2002), where its expression is regulated by ethylene, as demonstrated in multiple studies (Wakasa *et al*., 2006; Costa *et al*., 2011; Busatto *et al*., 2016). In particular, a specific allele of the gene *MdPG1* (*MDP0000326734*), *Md-PG1*_*SNP*_*-G*, is associated with medium/low texture and mealiness typical of ‘Golden Delicious’ (Costa *et al*., 2010; Longhi *et al*., 2013). In the first version of the apple genome (Velasco *et al*., 2010), this gene was located on chromosome 10 (chr10:18137615..18140009). Interestingly, the *PG* gene among the observed ASEs showing the highest value of baseMean (1035), namely GDH2-10g24673, is also located in the same region of chromosome 10 (Figure 3) (chr10_Hap2: 29418134..29420530) and belongs to haplotype H2. Furthermore, the identity between the two sequences is nearly complete (99%, with no gaps), supporting the hypothesis that GDH2-10g24673 corresponds to *Md-PG1*_*SNP*_*-G*. Conversely, the other allele *Md-PG1*_*SNP*_*-T* corresponds to GDH1-10g24691 and is positioned on chr10_Hap1: 29414507..29416903. Between the two, GDH2-10g24673 is upregulated compared to GDH1-10g24691, showing a log□fold change of 1.35 (padj = 2.15 × 10□^11^) and average allele specific raw counts of 1,607 and 591, respectively. Considering the higher expression levels of *Md-PG1*_*SNP*_*-G* (GDH2-10g24673) compared to *Md-PG1*_*SNP*_*-T* (GDH1-10g24691), it can be concluded that the association of *Md-PG1*_*SNP*_*-G* with increased flesh softening is due not only to intrinsic biochemical changes in the PG1 enzyme, resulting from three non-synonymous substitutions among the 18 SNPs distinguishing the two alleles (Longhi *et al*., 2013), but also its greater abundance.

**Figure 3.**
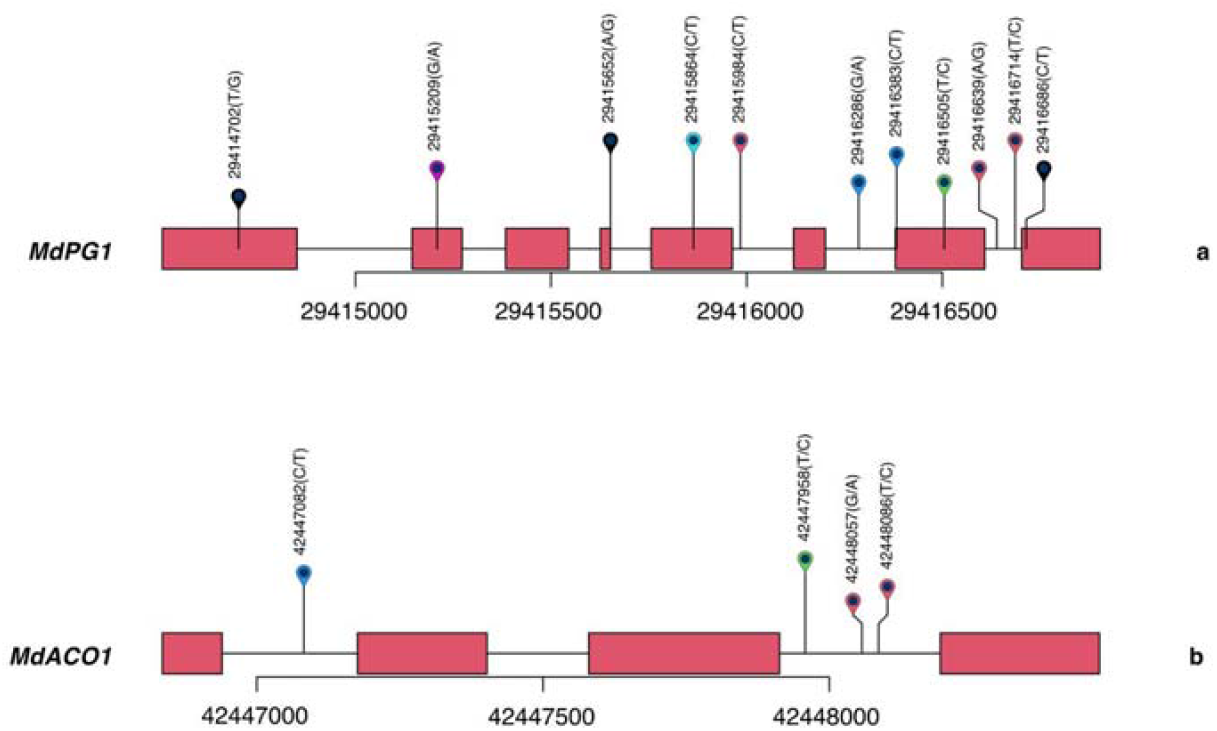
Distribution of SNPs in the genomic regions and gene structure of *MdPG1* (a) and *MdACO1* (b). Genomic coordinates on chr 10 are indicated along the x-axis, with each vertical line representing a single nucleotide variant distinguishing the two haplotypes. These SNPs are located within or near the coding sequences of *MdACO1* and *MdPG1*, two genes implicated in fruit ripening and texture in apple.

Therefore, it is possible to hypothesize that the specific softening behavior of ‘Golden Delicious’ is highly influenced by Haplotype2, which is likely inherited from the paternal lineage of unknown origin.

As a complementary objective, the role of ASE to the overall differential expression of transcripts detected by conventional analysis was examined. To this end, the two haplotypes were considered separately, each serving as the representative genome. For each haplotype, a set of differentially expressed genes (data from accessions PRJNA1399370) (DEGs) was identified by applying the thresholds *padj* ≤ 0.01 and |log□FC| ≥ 2, resulting in 109 DEGs for Haplotype1 and 111 for Haplotype2 between the two developmental stages considered (Supplementary Table S3 and S4). Among these DEGs, five allelic pairs were found in which one allele was expressed at least four-fold higher than its counterpart. Additionally, a case was observed involving two allelic (paralogous) pairs located on different chromosomes, where only one pair was expressed (Supplementary Table S5, S6 and Supplementary Table S7). Notably, in conventional DEG analysis, even non-expressed pairs are still reported as differentially expressed (Supplementary Table S7).

### Limitations in ASE quantification for loci with perfect sequence identity and resolution of contamination in homologous regions for allele prediction

Despite this type of analysis represents a significant step forward in explaining the molecular mechanisms contributing to the dominance of a single haplotype in specific traits (Minamikawa *et al*., 2021) and highlights the potential applications of haplotypes in crop breeding (Bhat *et al*., 2021) in other cases this approach can be less effective.

For example, this phenomenon is evident when studying one of the key genes responsible for climacteric ethylene production in apple: *1-aminocyclopropane-1-carboxylic acid oxidase 1* (*MdACO1*). This gene has been extensively studied due to its association with varying ripening patterns across different cultivars (Costa *et al*., 2005; Zhu *et al*., 2008). *MdACO1* catalyzes the final step in ethylene biosynthesis by converting ACC (1-aminocyclopropane-1-carboxylic acid) into ethylene. In fruit-bearing species such as apple, the expression of *MdACO1* is closely linked to the initiation and progression of fruit ripening, making it a key focus in postharvest physiology and shelf-life research (Houben and Van de Poel, 2019).

Costa *et al*. (2005, 2014) mapped *MdACO1* to the distal end of chromosome 10 (MDP0000195885, chr10:36882773–36884410), identifying two alleles originally distinguished by a large 62 bp indel located in the third intron, but not by any differences in the coding sequence. In this updated phased version of the genome, two collinear genes (Supplementary Table S8) with identical sequences (100% identity, no gaps) were identified on the two haplotypes on chromosome 10: *GDH1-10g25959* (chr10_Hap1: 42,446,835–42,448,472) and *GDH2-10g25947* (chr10_Hap2: 42,687,996–42,689,633). In this case, the differential expression analysis between the two alleles did not return statistically significant results (adjusted p-value = 0.18). This outcome is largely caused by the limitations of current RNA-seq mapping algorithms, which are unable to resolve read assignments between loci with 100% sequence identity on the two haplotypes, thus compromising accurate quantification of ASE.

Moreover, given that the two *ACO* alleles differ only in the size of the third intron, their discrimination is possible exclusively through analysis of the corresponding genomic sequence. To date, two conflicting reports exist in the literature regarding the allelic state of *MdACO1*. Zhu and Barritt (2008), based on the screening of 60 apple cultivars, identified ‘Golden Delicious’ as heterozygous (*MdACO1*-1/2), whereas Costa *et al*. (2014) reported the presence of only the *MdACO1*-2 allele. However, the alignment of the genomic sequences of *GDH1-10g25959* and *GDH2-10g25947* did not reveal the presence of this indel, showing instead only minimal differences: a single SNP in the first intron and three SNPs in the third intron (Figure 3B). Therefore, this updated version of the genome enabled to confirm the allelic state of *MdACO1* reported by Costa *et al*. (2014), with ‘Golden Delicious’ carrying only the *MdACO1*-2 allele.

### Allele-specific miRNA analysis

Using the full set of known apple miRNA, the mature and hairpin sequences were aligned against the two haplotypes to successfully annotate the known miRNA loci. One aspect highlighted by the analysis was that several miRNA members belonging to the multilocus families, were characterized by perfectly overlapping alignments on the same locus, yielding identical BLASTn results. Notably, the *M*.*domestica* miRNA hairpin annotations in miRBase refer to the contigs of the apple genome version 1.0. This has generated a redundant annotation, which is now evident with the availability of the 2 resolved haplotypes. In Supplementary Table S9 the miRNAs were filtered and a non-redundant list is provided. Differences were observed in the chromosomal distribution of the loci, since several families with numerous loci did not show a complete synteny. As examples, the miR171d was found only on chromosome 5 of Haplotype1 and was entirely absent from Haplotype2. However, in terms of function and regulatory capability associated with the miR171 family there should not be differences among the two haplotypes considering that all the other loci are conserved in both Haplotype1 and Haplotype2. Another miRNA, miR10981, was characterized by the presence of miR10981a on chromosome 10 of Haplotype1 only and miR10981c and d on chromosome 8 of Haplotype2 only. However, miR10981b being 100% identical to miR10981a in its mature form, and present in both Haplotype1 and Haplotype2 could compensate for the absence of miR10981a in Haplotype2. On the contrary, miR10981c and d are identical in their mature sequence but different compared to miR10981a and b. This could result in a different functional regulation of the miR10981c and d targets in a crossing population derived from ‘Golden Delicious’. MiR10981a-d were predicted in silico to target Apple Chlorotic Leaf Stem Virus (ACLSV) ORF1 (Ashraf *et al*., 2023) thus the overall putative function would be still provided by miR10981a and b in both Haplotype1 and Haplotype2. Similarly, miR10989 also targeting ACSLV was distributed non-synthenically with miR10989a present on Haplotype2 only and miR10989b, c, d, e on Haplotype1 only. Another miRNA, mdm-miR10993 showed differences between haplotypes where mdm-miR10993c, d and e were missing in Haplotype2. Like miR10989, miR10993 was predicted to target ACLSV in its c, d, e and f versions thus the regulatory capability could be conserved in both haplotypes through mdm-miR10993f. For mdm-miR11009 although conserved in both Haplotype1 and Haplotype2, its location for Haplotype1 was in chromosome 15 and in chromosome 14 in Haplotype2. An interesting case was represented by the miR7121 family, which is entirely present in Haplotype1 whereas only miR7121c was identified in Haplotype2 with all other family members appearing truncated. Generally, the differences observed between haplotypes for the aforementioned miRNAs would not result in functional divergence at the post-transcriptional level in a scenario where the two haplotypes segregate in a crossing population. However, other miRNAs were found to be specific to either one or the other haplotype Specifically, miR11012 and miR11016 (also potentially involved in the defence against ACSLV, Ashraf *et al*., 2023), along with miR10983 and miR10987 (both with unknown potential targets), were found only in Haplotype1. On the contrary, miR7128 was found exclusively in Haplotype2, but no targets are known for this miRNA either.

### Allele-specific methylation analysis

To investigate haplotype specific methylation, the ‘Golden Delicious’ apple genome was sequenced using Oxford Nanopore Technologies (ONT), yielding 42.13 Gb of high-quality data (reads N50 of 19.7 Kb and an average quality score >Q20). The 4.3 million reads were aligned to the concatenation of Haplotypes 1 and 2, resulting in an average raw read depth of 30x -35x on each haplotype, which decreased to 22.7x and 23.2x for Haplotype1 and 2, respectively, after removing the reads with a mapping quality lower than 10 and retaining only the primary alignments. The high number of reads with a mapping quality of 0 was due to identical loci in Haplotype 1 and Haplotype 2. The analysis of the combined haplotypes revealed that 20.5% of the cytosines are modified as 5-methylcytosine (5mC) and 0.21% as 5-Hydroxymethylcytosine (5hmC), with methylation levels varying significantly across chromosomes. The low abundance of 5hmC, consistent with observations in other plant species, appears to be a common feature of plant methylomes (Kumar *et al*., 2018). Specifically, Chromosome 1 of Haplotype 1 (chr1_Hap1) exhibited the highest 5mC percentage (24.1%), while Chromosome 9 of Haplotype 2 (chr9_Hap2) was the least methylated (18.8%).

The average methylation rates observed across the different sequence contexts were 85.0% for mCG, 65.7% for mCHG, and 5.9% for mCHH. These values are consistent with patterns reported in other fruit tree species, where CG and CHG contexts typically show high levels of methylation, while CHH sites display markedly lower frequencies. The overall methylation levels observed here are comparable to those reported for maize (Bartels *et al*., 2018).

The differential methylation between the 20,057 collinear genes present in both haplotypes in a single copy showed a highly similar methylation profiles, with only 44 genes that showed an absolute value of –Log2FoldChange above 2 (Supplementary Figure S4). As an example the differential methylation in the collinearity region between GDH2-10g25497 and GDH1-10g25500 was reported in Supplementary Figure S5. Even though this was only a feasibility study since only one physiological condition was analyzed, the use of long ONT reads, mapped to the two fully resolved ‘Golden Delicious’ haplotypes, proved instrumental in distinguishing their epigenetic landscapes. This approach successfully revealed distinct haplotype-specific methylation profiles, providing a proof-of-concept for exploring allele-specific epigenetic regulation in apple. Additionally, both haplotypes exhibited a higher methylation level in regions containing transposable elements (TEs) identified by RepeatMasker (see Material and Methods) than in collinear genes. In particular, the average methylation rates, in TE longer than 500nt, observed across the different sequence contexts were 97.5% for mCG, 90.7% for mCHG, and 24.0% for mCHH. As illustrated in the Supplementary Figure S6 and Table S10, 50% of TEs were located in regions with complete methylation (1.0 bin), and an additional 28% were found in the 90-99% methylation range (0.9 bin).

A comprehensive comparison is reported in Figure 4, which includes the TE signals, genes and methylation densities.

**Figure 4.**
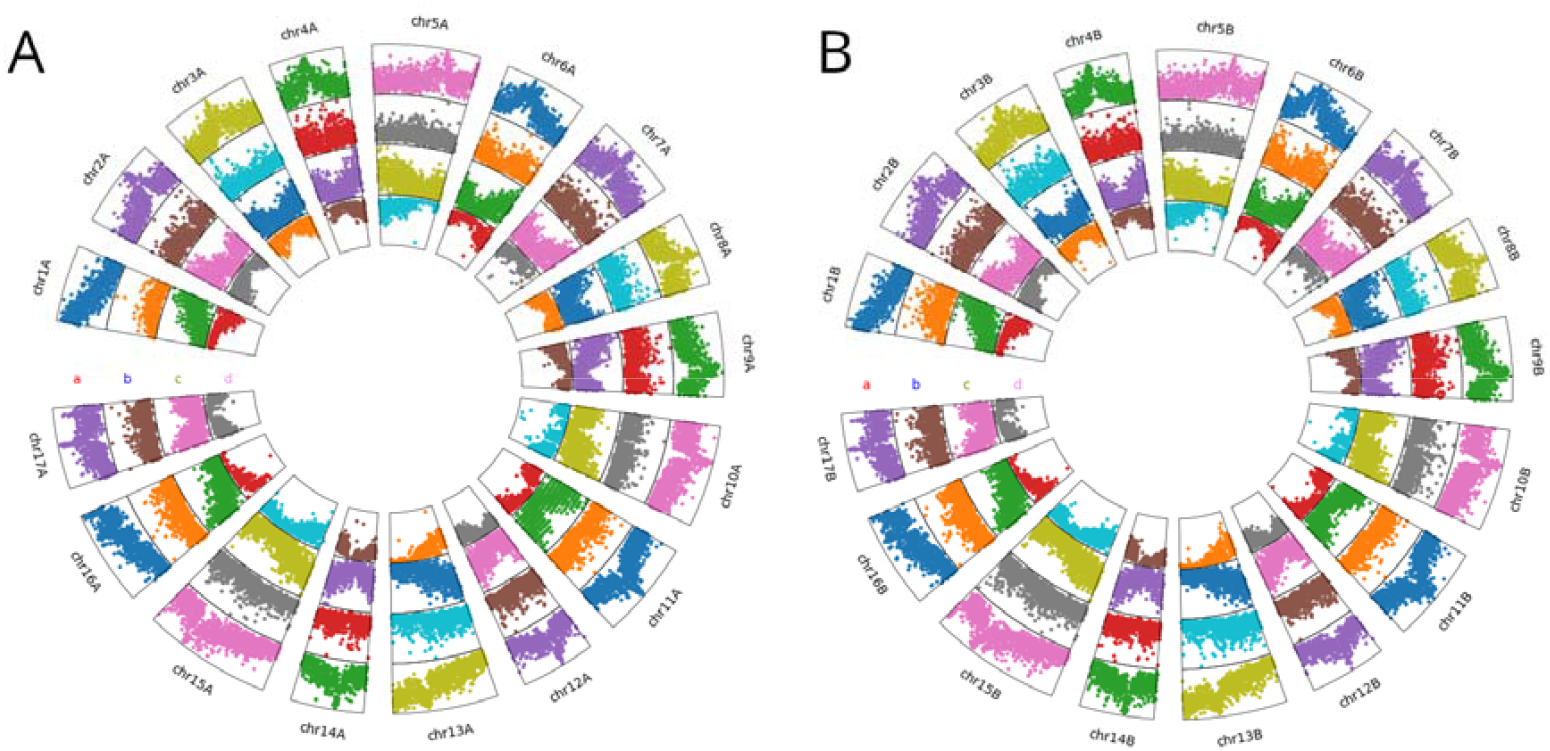
Circos plots for Golden Delicious’ Haplotype 1 (A) and Haplotype 2 (B). a) ClassI TEs, b) ClassII TEs, c) Gene density, d) CG methylation density.

## Conclusions

Haplotype-resolved genomes represent a paradigm shift in genetic analysis, moving beyond the limitations of the traditional collapsed consensus sequence. By providing a high-resolution, haplotype-specific reconstruction of parental alleles, this approach can enable the accurate identification of heterozygous regions and the precise analysis of both genetic and epigenetic variation within target genes. These foundational genomic resources are essential for understanding the allelic-specific contribution to plant phenotype, thereby significantly enhancing genomic selection and gene editing applications through improved target design and prediction of breeding outcomes. Future progress will rely on leveraging this haplotype-level resolution to dissect allelic-specific expression (ASE), which is fundamental to identifying the causal alleles and haplotype regions that must be maintained in breeding progeny. Furthermore, short-read Illumina RNA-seq i remains suboptimal for robust ASE analysis, particularly in genomes with extensive gene duplication (such as in apple), where read assignment is frequently problematic. Long-read sequencing technologies offer a promising alternative, by providing full-length transcript coverage, including the more variable 3’ sequence regions, which can dramatically improve allele-specific read assignment, paving the way for truly comprehensive, allele-specific transcriptome profiling.

## SUPPLEMENTARY DATA

The following supplementary data are available at JXB online.

**Supplementary Figure S1** Marey plots of Chr1-17 for ‘Golden Delicious’s’ Haplotype 1 and Haplotype 2.

**Supplementary Figure S2** Double inversion in Golden Delicious Haplotype 1 vs Haplotype 2, which is not present in Gala.

**Supplementary Figure S3** Allele-Specific Expression (ASE) landscape across the apple 17chromosomes.

**Supplementary Table S1** Genes in region 10.9M-16.4M on Haplotype 1 of chromosome

**Supplementary Table S2** Genes in region 13.2M-16.4M on Haplotype 2 of chromosome 6

**Supplementary Table S3** DESeq2 results of the differential expression analysis between the two time points using a count table obtained with a classic alignment approach on Haplotype1

**Supplementary Table S4** DESeq2 results of the differential expression analysis between the two time-points using a count table obtained with a classic alignment approach on Haplotype2

**Supplementary Table S5** DESeq2 results of the differential expression analysis between the two time points using a count table obtained considering the allele-specific expression data on Haplotype1 as input.

**Supplementary Table S6** DESeq2 results of the differential expression analysis between the two time points using a count table obtained considering the allele-specific expression data on Haplotype1 as input.

**Supplementary Table S7** Transcripts showing differential expression using the conventional approach with DESeq2.

**Supplementary Table S8** gene collinearity between the two haplotypes.

**Supplementary Table S9** miRNA families annotation table with coordinates on Haplotype1 and Haplotype2.

**Supplementary table S10** Absolute counts of TE elements per methylation bin gene collinearity between the two haplotypes.

## Acknowledgements

Authors would like to thank Nicholas Howard, Azeddine Si Ammour, Alessandro Botton, Francesca Populin, and Elisa Asquini for providing access to data used in this work.

## Author contributions

LB, NB, MM, DM, MT, FC, PF: Conceptualization; LB, PF: Data Curation; LB, NB, MM, DM, LS, PF: Formal Analysis; MM, SP: Investigation; MT: Resources; LB, NB, MM, DM, LS, MT, SP, FC, PF: Writing - Original Draft Preparation and Review & Editing;

## Data availability

The assembled sequence in.fasta format and gene predictions in gff3 format are publicly available through the Genome Database for Rosaceae (GDR) at the following URL: www.rosaceae.org/Analysis/27106201

PacBio and ONT raw reads are available under the Bioproject id PRJNA1402909. The gene functional annotation is publically available at https://doi.org/10.6084/m9.figshare.31123414

